# Shared and disease-specific host gene-microbiome interactions across human diseases

**DOI:** 10.1101/2021.03.29.437589

**Authors:** Sambhawa Priya, Michael B. Burns, Tonya Ward, Ruben A. T. Mars, Beth Adamowicz, Eric F. Lock, Purna C. Kashyap, Dan Knights, Ran Blekhman

## Abstract

While the gut microbiome and host gene regulation separately contribute to gastrointestinal disorders, it is unclear how the two may interact to influence host pathophysiology. Here, we developed a machine learning-based framework to jointly analyze host transcriptomic and microbiome profiles from 416 colonic mucosal samples of patients with colorectal cancer, inflammatory bowel disease, and irritable bowel syndrome. We identified potential interactions between gut microbes and host genes that are disease-specific, as well as interactions that are shared across the three diseases, involving host genes and gut microbes previously implicated in gastrointestinal inflammation, gut barrier protection, energy metabolism, and tumorigenesis. In addition, we found that mucosal gut microbes that have been associated with all three diseases, such as *Streptococcus*, interact with different host pathways in each disease, suggesting that similar microbes can affect host pathophysiology in a disease-specific manner through regulation of different host genes.

## Introduction

The human gut microbiome plays a critical role in modulating human health and disease. Variations in the composition of the human gut microbiome have been associated with a wide variety of chronic diseases, including colorectal cancer (CRC), inflammatory bowel disease (IBD), and irritable bowel syndrome (IBS). For example, previous studies have reported an increase in abundance of *Fusobacterium nucleatum* and *Parvimonas* in CRC ^1,2^, reduced abundance of *Faecalibacterium prausnitzii* and enrichment of enterotoxigenic *Bacteroides fragilis* in CRC and IBD ^3–5^, and overrepresentation of Enterobacteriaceae and *Streptococcus* in IBD and IBS ^6–8^. In addition to the gut microbiome, dysregulation of host gene expression and pathways have also been implicated in these diseases. Researchers have reported disruption of Notch and WNT signalling pathways in CRC ^9,10^, activation of toll-like receptors (e.g. TLR4) that induce NF-κB and TNF-α signaling pathways in IBD ^11,12^, and dysregulation of immune response and intestinal antibacterial gene expression in IBS ^8,13^. While host transcription and gut microbiome have separately been identified as contributing factors to these gastrointestinal (GI) diseases, it is unclear how the two may interact to influence host pathophysiology ^14^.

Studies in model organisms have demonstrated that the modulation of host gene expression by the gut microbiome is a potential mechanism by which microbes can affect host physiology ^15–20^. For example, in zebrafish, the gut microbiome negatively regulates the transcription factor hepatocyte nuclear factor 4, leading to host gene expression profiles associated with human IBD ^18^. In mice, the gut microbiota can alter host epigenetic programming to modulate intestinal gene expression involved in immune and metabolic processes ^16,17^. Additionally, recent *in vitro* cell culture experiments have shown that specific gut microbes can modify the gene expression in interacting human colonic epithelial cells ^21,22^. Given the evidence for crosstalk between the gut microbiome and host gene regulation, characterizing the interplay between the two factors is critical for unravelling their role in the pathogenesis of human intestinal diseases.

A few recent studies have investigated interactions between the host transcriptome and gut microbiome in specific human gut disorders, including IBD, CRC, and IBS. For example, studies examining microbiome-host gene relationships in IBD have identified mucosal microbiome associations with host transcripts enriched for immunoinflammatory pathways ^23–25^. While investigating longitudinal host-microbiome dynamics in IBD, Lloyd-Price et al. identified interactions between expression of chemokine genes, including *CXCL6* and *DUOX2*, and abundance of gut microbes, including *Streptococcus* and Ruminococcaceae ^25^. Studies investigating the role of host gene-microbiome interactions in CRC have found correlations between the abundance of pathogenic mucosal bacteria and expression of host genes implicated in gastrointestinal inflammation and tumorigenesis ^26,27^. In IBS, host genes implicated in gut barrier function and peptidoglycan binding, such as *KIFC3* and *PGLYRP1*, are associated with microbial abundance of Peptostreptococcaceae and *Intestinibacter* ^8^. While these studies have revealed important insights about host gene-microbiome crosstalk in GI diseases, they are limited in several aspects. For example, to boost statistical power, most studies have examined interactions between a limited subset of host genes and gut microbes; for instance, by focusing only on differentially expressed genes ^24,25,27^, genes associated with immune functions ^13,26^, or select microbes representing bacterial clusters or co-abundance groups ^23,26^, thus characterizing only a subset of potential interactions. In addition, the identification of host gene-microbe interactions is based on testing for pairwise correlation between every host gene and microbe using Spearman or Pearson correlation, thus ignoring the inherent multivariate properties of these datasets ^24,25,27^. This approach may also decrease statistical power to detect biologically meaningful associations due to the large number of statistical tests performed. Additionally, most studies focus on examining interactions in a single disease at a time; hence, common and unique patterns of host-microbiome interactions across multiple disease states remain poorly characterized.

Here, we comprehensively characterized interactions between mucosal gene expression and microbiome composition in patients with colorectal cancer, inflammatory bowel disease, and irritable bowel syndrome, three GI disorders in which both host gene regulation and gut microbiome have been implicated as contributing factors ^1,6,8,10,13^. We developed and applied a machine learning framework that overcomes typical challenges in multi-omic integrations, including high-dimensionality, sparsity, and multicollinearity, to identify biologically meaningful associations between gut microbes and host genes and pathways in each disease. We leveraged our framework to characterize disease-specific and shared host gene-microbiome interactions across the three diseases that may facilitate new insights into the molecular mechanisms underlying pathophysiology of these gastrointestinal diseases.

## Results

### Integrating host gene expression and gut microbiome abundance in colorectal cancer, inflammatory bowel disease and irritable bowel syndrome

To study host-microbiome relationship across diseases, we used host gene expression (RNA-seq) data and gut microbiome abundance (16S rRNA sequencing) data generated from colonic mucosal biopsies obtained from patients with colorectal cancer (CRC), inflammatory bowel disease (IBD), and irritable bowel syndrome (IBS) (**Figure 1A**). Our study included 208 microbiome samples and 208 paired gene expression samples (416 in total). These 208 paired samples include 88 pairs of samples in the CRC cohort (44 tumor and 44 patient-matched normal), 78 pairs of samples in the IBD cohort (56 patients and 22 controls)^25–28^, and 42 pairs of samples in the IBS cohort (29 patients and 13 controls; see Supplementary Table S1)^8^. Detailed information on disease cohorts, samples, sequencing, quality control, and data processing is available in Methods.

**Figure 1.**
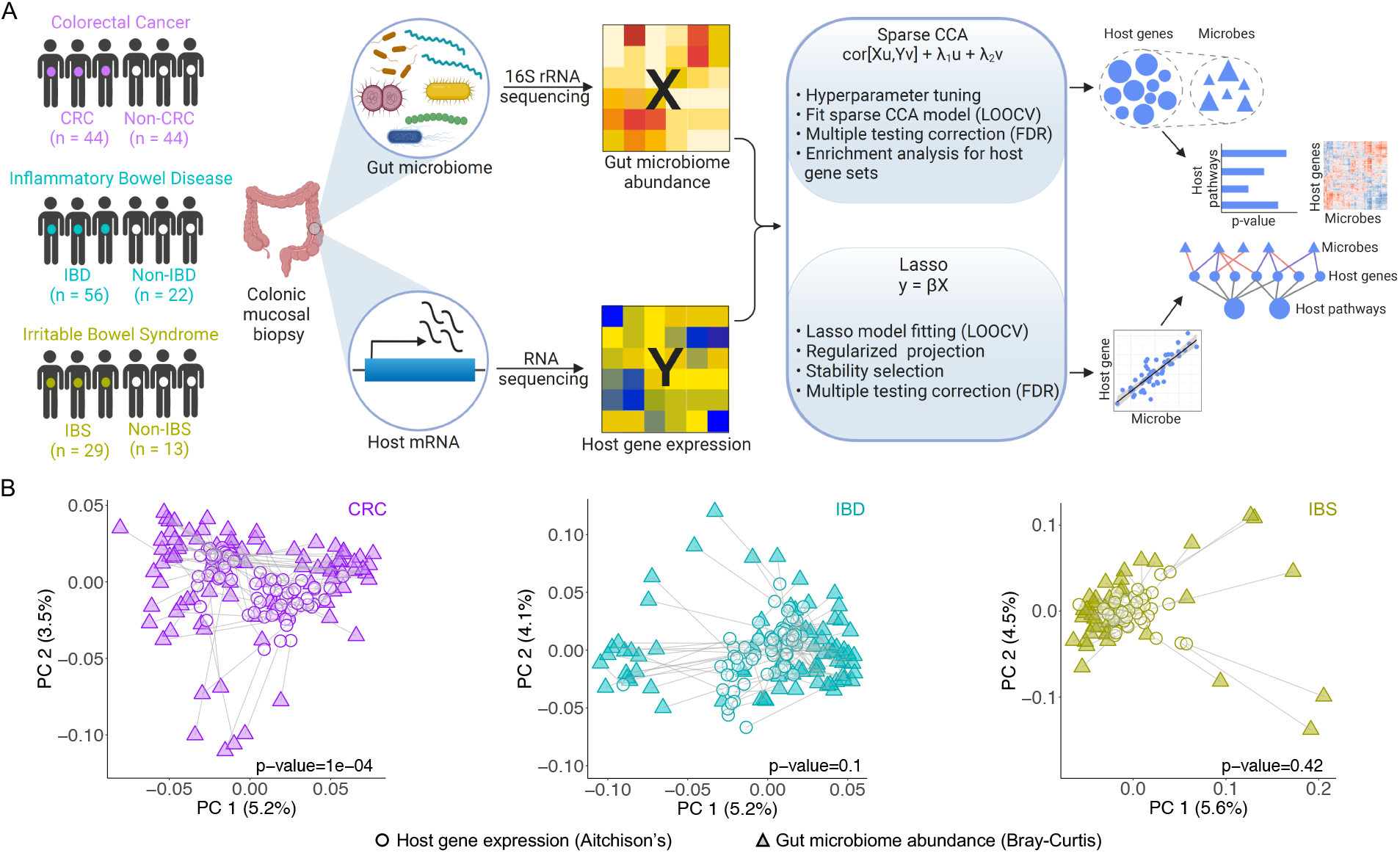
Integrating host gene expression and gut microbiome abundance in colorectal cancer (CRC), inflammatory bowel disease (IBD), and irritable bowel syndrome (IBS). **A.** Study design representing disease cohorts, generation of host gene expression data and gut microbiome abundance data from patient samples, overview of integration framework, and expected output (left to right). For description of mathematical notations, please see Methods. **B.** Procrustes analysis showing overall association between variation in host gene expression and gut microbiome composition in CRC, IBD and IBS (left to right). We used Aitchison’s distance for host gene expression data (circle), and Bray-Curtis distance for gut microbiome data (triangle).

Previous studies have identified host gene-microbiome associations in human gut disorders, including CRC, IBD, and IBS^8,25,26^. Thus, one might expect intestinal gene expression patterns and microbiome composition to be broadly correlated in these diseases. To test for such an overall association between host gene expression and gut microbiome composition, we performed Procrustes analysis using paired data for each disease cohort. Our analysis showed significant correspondence between host gene expression variation and gut microbiome composition across subjects in CRC (Monte Carlo p-value = 0.0001). However, Procrustes agreement is not significant in IBD (Monte Carlo p-value = 0.1) and IBS (Monte Carlo p-value = 0.42) (**Figure 1B**, see Methods). This lack of significant overall correspondence between host transcriptome and gut microbiome across diseases might suggest that, instead of an overall association between the two, it is likely that only a subset of gut microbes interact with a subset of host genes at the colonic epithelium ^15,17^. Hence, we need novel integration approaches to characterize such host gene-microbiome interactions.

To this end, we developed a machine learning framework for integrating multi-omic high-dimensional datasets, such as host gene expression and gut microbiome abundance, to identify relevant host genes and pathways associated with gut microbes. Our integration approach has two parts: (i) Sparse canonical correlation analysis (sparse CCA) ^29,30^ for identifying groups of host genes that associate with groups of gut microbial taxa to characterize pathway-level interactions; and (ii) Lasso penalized regression ^31^, for identifying specific interactions between individual host genes and gut microbial taxa (see **Figure 1**, Methods, Supplementary Figure S1). We applied our integration analysis to matched host gene expression data and gut microbiome data for each disease cohort separately to avoid any potential batch effects. For each disease cohort dataset, we conducted the integration analysis separately for the patient data (i.e. CRC, IBD, and IBS) and corresponding control data (non-CRC, non-IBD, and nonIBS, respectively), and considered only associations that were found in patients and not in controls. As opposed to the Procrustes analysis, our approach identified significant and potentially biologically meaningful associations between gut microbiota and host genes and pathways across the three diseases.

### Shared host pathways associate with disease-specific gut microbes across GI diseases

We hypothesized that host genes and gut microbial taxa involved in common biological functions would act in a coordinated fashion, and, hence, would have correlated expression and abundance patterns. To investigate this, we used sparse CCA to characterize group-level association between host transcriptome and gut microbiome in each of the three diseases ^29,30^. We fit the sparse CCA model for each dataset to identify subsets of significantly correlated host genes and gut microbes, known as components (see Methods, and Supplementary Tables S2–S4). We then performed pathway enrichment analysis on the set of host genes in each significant component to determine host pathways that interact with gut microbes in a disease. We identified *shared* pathways, namely host pathways for which gene expression correlates with gut microbes across disease cohorts, and *disease-specific* pathways, namely host pathways for which gene expression correlates with gut microbes in only one of the three disease cohorts (**Figure 2A**; Fisher’s exact test, Benjamini-Hochberg FDR < 0.1, Supplementary Table S5). For simplicity, we focused on the top five most significant shared and disease-specific pathways (**Figure 2A**). We found three pathways shared across CRC, IBD, and IBS that are known to regulate gastrointestinal tract inflammation and gut barrier protection and repair. For example, oxidative phosphorylation, which is the process of energy metabolism in the mitochondria, is known to be upregulated in IBD and CRC, and contributes to tumorigenesis and drug resistance in CRC ^32–35^. Interestingly, the gut microbiome can signal mitochondria of gut mucosal immune cells to alter mitochondrial metabolism, including oxidative phosphorylation processes; this can lead to impaired epithelial barrier function and chronic intestinal inflammation in IBD and CRC ^36^. We also found overlapping host pathways between disease pairs (see CRC & IBD, CRC & IBS, and IBD & IBS in **Figure 2A**), including immunoregulatory pathways and cell-surface receptors like integrin pathway, cell and focal adhesion, and proteasome.

**Figure 2.**
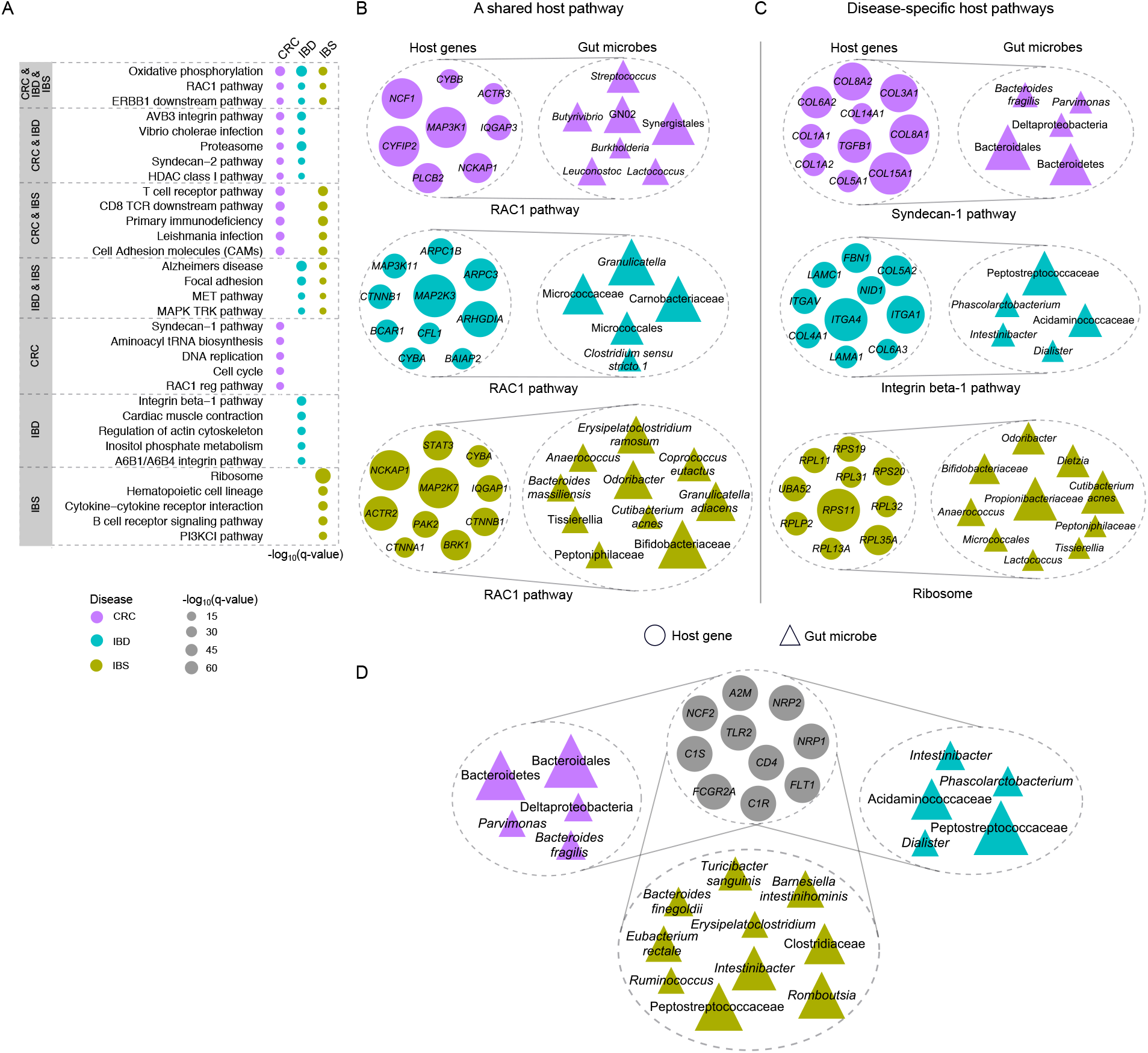
Shared immunoregulatory and metabolic host pathways associate with disease-specific gut microbes across human diseases. **A.** Host pathways enriched for sparse CCA gene sets associated with gut microbiome composition across diseases (FDR < 0.1). Size of the dots represent the significance of enrichment for each pathway, and color of the dots denote the disease cohort in which this pathway is significantly associated with microbiome composition. **B.** Association between microbial taxa in CRC, IBD and IBS (top to bottom) and host genes in the RAC1 pathway, a shared host pathway (i.e., a pathway for which host gene expression correlates with gut microbes across disease cohorts). Size of circles and triangles represent the absolute value of sparse CCA coefficients of genes and microbes, respectively. **C**. Association between the set of host genes in disease-specific host pathways (i.e. host pathways for which gene expression correlates with gut microbes in only one of the disease cohorts) and group of gut bacteria in CRC, IBD and IBS (top to bottom). **D**. A common set of host genes (grey circles) interact with disease-specific sets of microbes. These host genes are enriched for immunoregulatory and acute inflammatory response pathways

In addition, we identified 102 disease-specific host pathways that are associated with gut microbes, including 52 CRC-specific, 25 IBD-specific, and 25 IBS-specific pathways (Supplementary Table S5, **Figure 2A**). While IBD-specific host pathways include A6B1/A6B4 integrin pathway and Integrin beta-1 pathway that regulate leukocyte recruitment in GI inflammation ^37,38^, IBS-specific pathways include immune response pathways, including B cell receptor signaling pathway, and ribosome pathway.

To better understand the host gene-microbe interactions that underlie common associations, we focused on the RAC1 pathway, where host gene expression is associated with microbiome composition in CRC, IBD, and IBS. The RAC1 pathway is known to regulate immune response and intestinal mucosal repair, and has previously been implicated in IBD and CRC ^39–41^ (**Figure 2B**). As expected, we observed overlapping host genes for this shared pathway across the three diseases. However, the microbial taxa they are correlated with are disease-specific. In CRC, the RAC1 pathway is associated with oral bacterial taxa such as *Streptococcus, Synergistales*, and *GN02*, where *Streptococcus* species are known to be associated with colorectal carcinogenesis ^42,43^. In IBD, the RAC1 host pathway is associated with microbial taxa previously implicated in IBD, including *Granulicatella ^44–46^*, and *Clostridium sensu stricto 1*, a microbe associated with chronic enteropathy similar to IBD ^47^. In IBS, this pathway is associated with bacteria such as *Bacteroides massiliensis*, that has been shown to be prevalent in colitis ^48^, and *Bifidobacterium* and *Odoribacter*, that are known to be depleted in IBS ^49–52^.

To investigate disease-specific associations, we considered unique host pathways for which host gene expression correlates with gut microbes only in one of the three diseases (**Figure 2C**). For example, the Syndecan-1 pathway, which we found to be associated with gut microbial taxa only in CRC, has been previously shown to regulate the tumorigenic activity of cancer cells by altering extracellular matrix adhesion and cell morphology ^53–55^. Host gene expression in this pathway is associated with microbial taxa such as *Parvimonas* and *Bacteroides fragilis* that are known to promote intestinal carcinogenesis and are considered biomarkers of CRC ^1, 56–58^. The integrin-1 pathway, a disease-specific host pathway in IBD, is found to be associated with *Peptostreptococcaceae, Intestinibacter*, and *Phascolarctobacterium*, microbial taxa that are have been implicated in IBD by previous studies ^59–62^. To assess similarities in host gene components across diseases, we identified a set of host genes that are common between components across the three diseases, and we found that these genes are enriched for immune response pathways in gut epithelium, including vascular endothelial growth factor (VEGF), complementation and coagulation cascades, and cytokine-cytokine receptor interaction (**Figure 2D;** Fisher’s exact test, Benjamini-Hochberg FDR < 0.1). While this set of host genes is associated with disease-specific groups of microbes, we also found overlapping microbes between IBD and IBS, such as *Peptostreptococcaceae* and *Intestinibacter*, taxa that are found in high abundance in gastrointestinal inflammation ^59,61,62^.

### Specific gut microbes interact with individual host genes and pathways in each disease

Previous studies have shown that specific microbial taxa can regulate expression of individual host genes ^15,22^. Therefore, we explored interactions between individual host genes and gut microbes in each disease. To do so, we used Lasso penalized regression models to identify specific gut microbial taxa whose abundance is associated with the expression of a host gene ^31^. We fit these models in a gene-wise manner, using expression for each host gene as response and abundance of gut microbial taxa as predictors. We then applied stability selection to identify robust associations (see Methods). Using this approach, we found 755, 1295, and 441 significant and stability-selected host gene-taxa associations in CRC, IBD, and IBS, respectively (**Figure 3;** Tables S6-S8; FDR < 0.1). These represent interactions between 745 host genes and 120 gut microbes in CRC (Supplementary Table S6), between 1246 host genes and 56 gut microbes in IBD (Supplementary Table S7), and between 436 host genes and 102 gut microbes in IBS (Supplementary Table S8) (**Figure 3A**). Examples of specific host gene-microbe interactions can be found in Supplementary Figure S2. Overall, we observed disease-specific patterns in host gene-taxa interactions.

**Figure 3.**
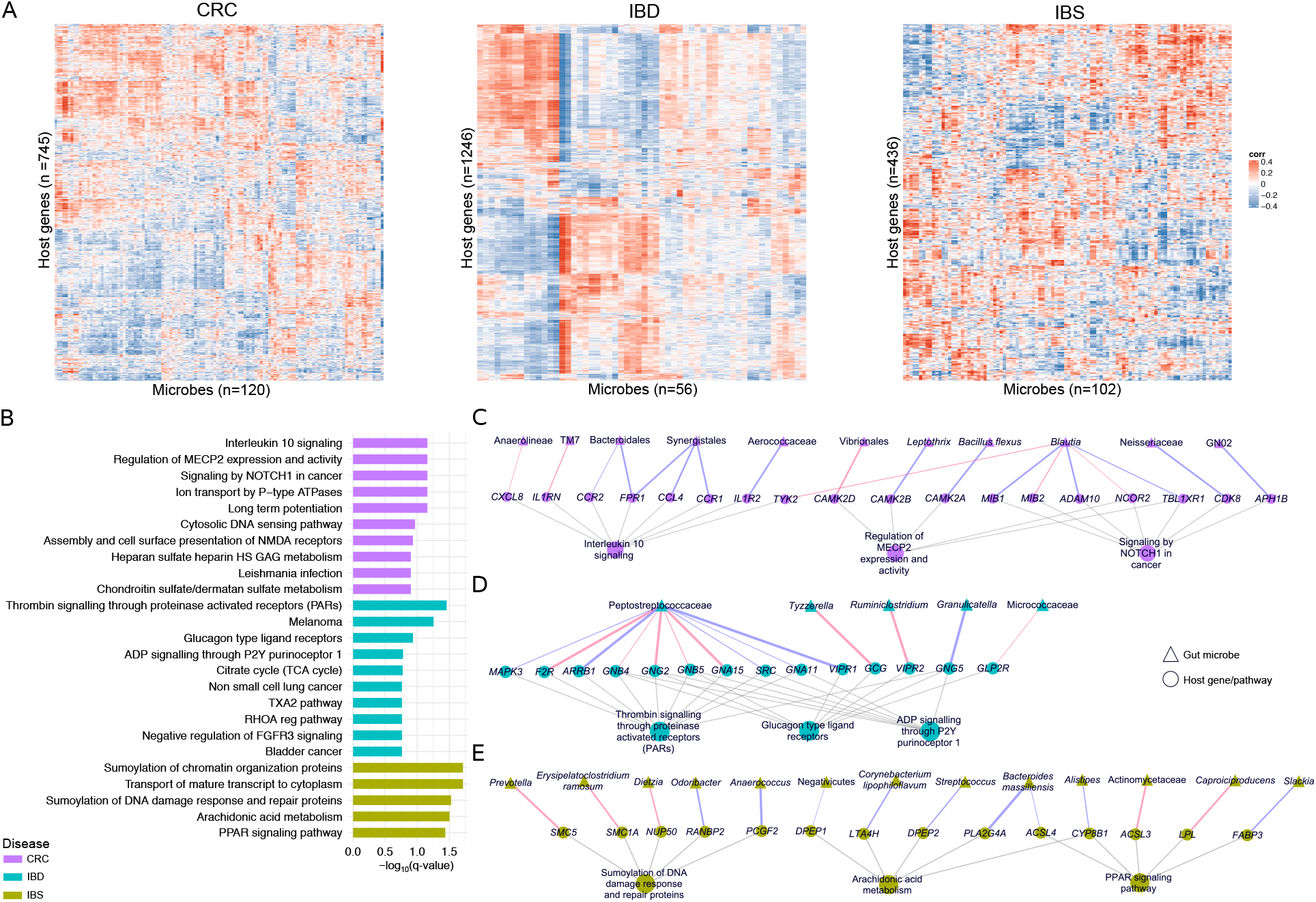
Specific gut microbes interact with individual host genes and pathways in each disease. **A.** Heatmap showing the overall pattern of interactions between significant and stability-selected host genes and gut microbial taxa identified by the Lasso model in CRC, IBD, and IBS (FDR < 0.1). **B.** Host pathways enriched among genes that are correlated with specific gut microbes in CRC (purple), IBD (green), and IBS (yellow). **C-D.** Networks showing specific gut microbes correlated with specific host genes enriched for disease-specific host pathways in CRC (**C**), IBD (**D**), and IBS (**E**). Triangular nodes represent gut microbes, circular nodes represent host genes and pathways. Edge color represents positive (blue) or negative (red) association, and edge width represents strength of association (spearman rho). Grey edges represent host gene - pathway associations.

To characterize the biological functions represented by the host genes that interact with specific gut microbes, we applied enrichment analysis on the set of gut microbiota-associated host genes in each disease (see Methods). This is complementary to our group-level approach (Figure 2) in that these host pathways are enriched among individual host gene-microbe pairs. We identified 87 host pathways that are unique to each disease, including 22 CRC-specific, 60 IBD-specific, and 5 IBS-specific pathways that interact with unique gut bacteria, of which we visualized top 10 most significant host pathways per disease (**Figure 3B**, Fisher’s exact test, Benjamini-Hochberg FDR < 0.2, Supplementary Table S9, see Methods). The host pathways enriched for CRC-specific interactions are known to modulate tumor growth, progression and metastasis in CRC, such as Interleukin-10 signaling, signaling by *NOTCH1* in cancer, and regulation of MECP2 expression and activity ^63–66^. The host pathways we identified as enriched for IBD-specific interactions are known to be responsible for maintenance of gastric mucosa integrity, inflammatory response, and host defence against invading pathogens, such as thrombin signalling through proteinase activated receptors (PARs), and glucagon type ligand receptors ^67,68^. For IBS-specific interactions, the enriched host pathways identified here have been shown to regulate homeostasis of intestinal tissue and proinflammatory mechanisms in IBS, such as sumoylation of DNA damage response and repair proteins, and arachidonic acid metabolism ^69–71^.

To characterize the potential mechanism of host gene-microbe interactions, we further investigated the gut microbial taxa associated with host genes in these pathways (**Figure 3C-E**). In CRC, we found that Anaerolineae and *TM7*, oral microbes that also inhabit the human gastrointestinal tract, and are known to promote oral and colorectal tumorigenesis ^72–76^, are negatively correlated with host genes enriched for tumor-promoting Interleukin-10 signaling pathway, such *CXCL8* and *IL1RN* (**Figure 3C** and Supplementary Figure S2). *CXCL8* is known to be overexpressed in CRC, and *IL1RN* is centrally involved in immune and inflammatory response, and its polymorphisms are implicated in colorectal carcinogenesis ^77–79^. Other host genes in Interleukin-10 signaling, such as *CCR2* and *FPR1*, are positively correlated with Bacteroidales (**Figure 3C** and Supplementary Figure S2). *CCR2* and *FPR1* are overexpressed in colorectal tumors, while Bacteroidales are enriched in CRC and associated with tumorigenesis ^80,82^

We observed that Peptostreptococcaceae, which is prevalent in patients with IBD ^62,83,84^, is associated with multiple host genes and pathways in IBD (**Figure 3D**). For example, its abundance is positively correlated with the expression of host genes *MAPK3* and *VIPR1*, involved in thrombin signalling through proteinase activated receptors (PARs) and glucagon type ligand receptors pathways, respectively. *MAPK3* is known to play a role in progression and development of IBD, and *VIPR1* is overexpressed in inflamed mucosa ^85,86^. The abundance of Micrococcaceae, which is known to be increased in IBD, is negatively associated with the expression of *GLP2R, a* glucagon receptor involved in maintenance of gut barrier integrity ^68,87^. In IBS-specific interactions, we found that the levels of Prevotella, which is known to be overrepresented in individuals with loose stool, to be negatively associated with expression of *SMC5*, which is involved in the sumoylation pathway ^88,89,88,90^ (**Figure 3E**). Previous studies have shown that gut pathogens can target the host sumoylation machinery that regulates inflammatory cascade in epithelial cells in inflammatory bowel disease ^69^. We also found the expression of *PLA2G4A*, a host gene that plays an important role in arachidonic acid metabolism and is an integral member of prostaglandin biosynthesis pathway that modulates gut epithelial homeostasis ^91,92^, is positively correlated with the abundance of *Bacteroides massiliensis* in IBS, a gut microbe known to be prevalent in patients with gut malignancies, including ulcerative colitis and colorectal carcinoma ^48,93^ (**Figure 3E**). Taken together, these findings demonstrate that interactions between specific gut microbial taxa and specific host genes and pathways vary by disease state.

### Disease-specific gut microbe-host gene crosstalk

To understand how shared gut microbes may interact with specific host genes across diseases, we explored the overlaps between host gene-microbe associations in CRC, IBD, and IBS (**Figure 4A**, Lasso regression, Benjamini-Hochberg FDR < 0.1, Supplementary Table S10). We found that the abundance of 3 gut microbes, Peptostreptococcaceae, *Streptococcus*, and *Staphylococcus*, is correlated with host gene expression in all three diseases (**Figure 4A; Network 1**). Previous studies have revealed that Peptostreptococcaceae and *Streptococcus* spp. are found at elevated levels in CRC, IBD, and IBS ^8,43,62,94,–97^. While traditionally considered nasal- or skin-associated bacteria, *Staphylococcus* spp. also colonize the human gastrointestinal tract and include opportunistic pathogens that can cause acute intestinal infections in patients with CRC and IBD ^98–103^, and are associated with increased risk of IBS and CRC ^97,103,104^. We found that the abundance of Peptostreptococcaceae is positively correlated with the expression of host genes *PYGB* and *NCK2* in IBD, whereas it is negatively correlated with the expression of host gene *HAS2* in IBS. *PYGB* and *NCK2* are both upregulated in IBD, where *PYGB* is known to regulate Wnt/p-catenin pathway, and *NCK2* is involved in integrin and epidermal growth factor receptor signaling ^105–109^. In contrast, *HAS2* is known to have a protective effect on the colonic epithelium through regulation of intestinal homeostasis and inflammation ^110–112^. In CRC, we found that the abundance of Peptostreptococcaceae is negatively associated with the expression of *GAB1*, a host gene for which overexpression stimulates tumor growth in colon cancer cells ^113^. *Streptococcus* also shows a disease-specific pattern of association with host gene expression. In CRC, its abundance is correlated with the expression of *RIPK4*, which regulates WNT signaling and NF-KB pathway, and is upregulated in several cancer types, including colon cancer ^114–116^. Similarly, in IBS, *Streptococcus* abundance is correlated with the expression of *DPEP2*, which is known to modulate macrophage inflammatory response ^117^, and is involved in arachidonic acid metabolism that is known to be dysregulated in IBS ^70,71^ (**Figure 4A; Network 1).**

**Figure 4.**
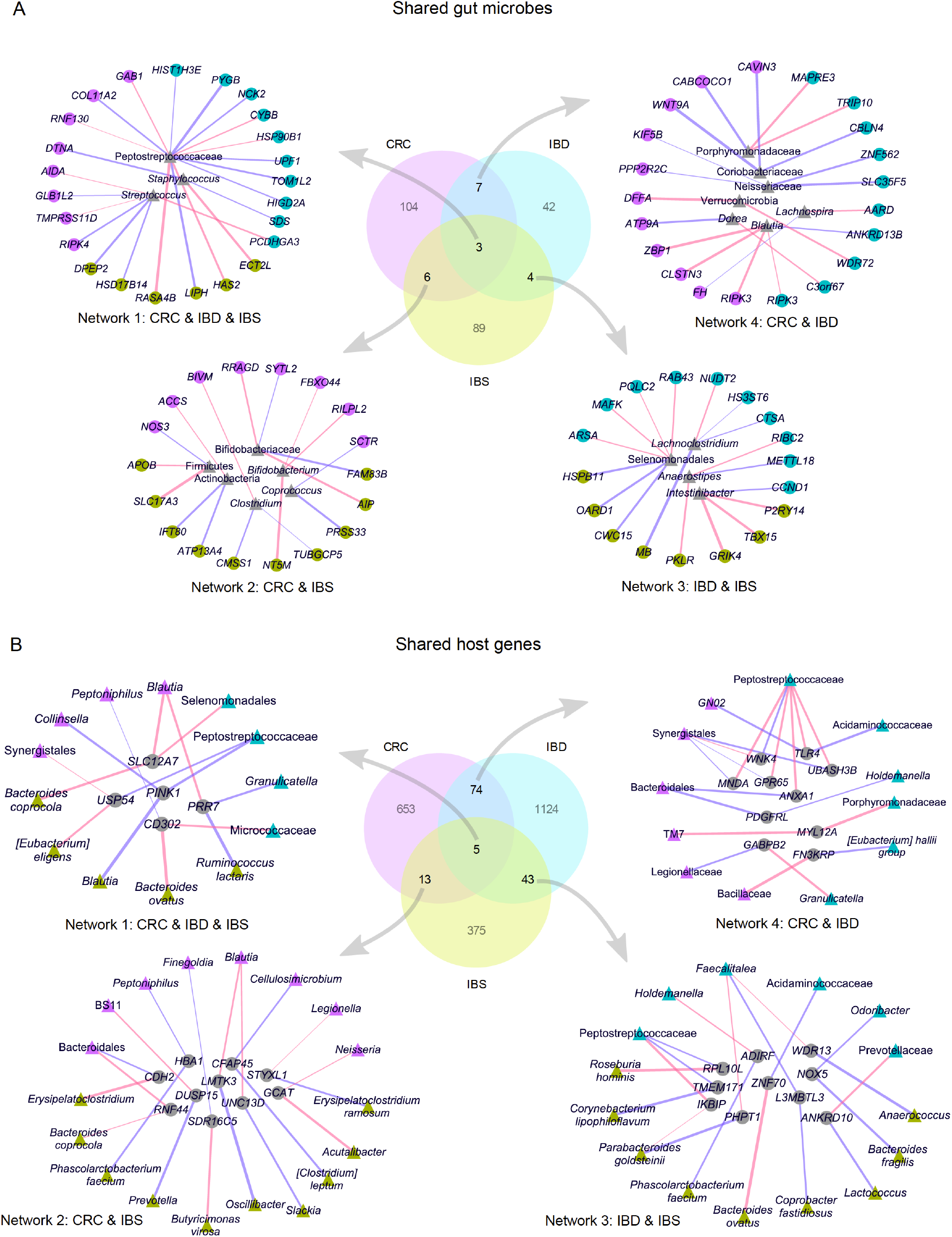
Associations for shared gut microbes and shared host genes reveal disease-specific host-microbiome crosstalk. **A.** *(center)* Venn diagram showing overlap between gut microbes associated with host genes in CRC, IBD and IBS, (*counter-clockwise*) networks showing host gene-microbe interactions for gut microbes shared across CRC, IBD and IBS (**Network 1),** between CRC and IBS (**Network 2**), between IBD and IBS (**Network 3**), and between CRC and IBD (**Network 4). B.** *(center)* Venn diagram showing overlap between host genes associated with gut microbes in CRC, IBD, and IBS, (*counter-clockwise*) networks showing host gene-microbe interactions for host genes shared across CRC, IBD and IBS (**Network 1**), between CRC and IBS (**Network 2**), between IBD and IBS (**Network 3**), and between CRC and IBD **(Network 4**). Circular nodes represent host genes, triangular nodes represent gut microbes. Colored nodes represent specific disease (purple: CRC, green: IBD, yellow: IBS), grey nodes represent gut microbes (A) and host genes (B) shared across diseases. Edge color represents positive (blue) or negative (red) association, and edge width represents strength of association (spearman rho). All interactions were determined at FDR < 0.1.

To elucidate potential host gene-microbe interactions for gut microbes shared between diseases, we visualized networks of most significant associations (**Figure 4A; Networks 2-4**, Lasso regression, Benjamini-Hochberg FDR < 0.1, Supplementary Table S10). We found 20 microbes for which abundance is associated with the expression of host gene in at least two diseases. Notably, the abundance of *Blautia, a* butyrate-producing beneficial microbe, is found to be negatively correlated with the expression of *RIPK3* in both CRC and IBD (**Figure 4A; Network 4** and Supplementary Figure S3). *RIPK3* promotes intestinal inflammation in IBD, and colon tumorigenesis ^118–122^. Interestingly, in CRC, *Blautia* is also associated with *ZBP1* (**Figure 4A; Network 4),** a host gene that recruits *RIPK3* to induce NF-KB activation, and regulates innate immune response to mediate host defense against tumors and pathogens ^123–125^.

Conversely, to explore how shared host genes may interact with gut microbes across all diseases, we identified host genes for which expression is correlated with the abundance of specific gut microbes in CRC, IBD, and IBS **(Figure 4B**, Lasso regression, FDR < 0.1, Supplementary Table S11). We identified 5 such host genes that interact with 4 gut microbes in CRC, 5 gut microbes in IBS, and 4 gut microbes in IBD **(Figure 4B; Network 1**, Supplementary Table S11). Of note, the expression of *PINK1*, a host gene that regulates mitochondrial homeostasis and activates PI3-kinase/AKT signaling, contributing to intestinal inflammation in IBD, and tumorigenesis ^126–128^, is associated with the abundance of *Collinsella* in CRC, Peptostreptococcaceae in IBD, and *Blautia* in IBS. Previous studies have found that *Collinsella* is increased in abundance in CRC, and has been shown to induce inflammation via altering gut permeability ^129–131^. However, *Blautia* has been found to be both positively and negatively correlated with IBS symptoms ^8,51,132^.

In addition, we identified 135 host genes for which expression is associated with abundance of microbial taxa in at least two of the three diseases, and visualized the network of most significant associations (**Figure 4B; Networks 2-4**, Lasso regression, FDR < 0.1, Supplementary Table S11). We found that the host genes whose expression is correlated with gut microbes in both CRC and IBD are enriched for pathways involved in immune response, including natural killer cell mediated toxicity, *Leishmania* infection, and leukocyte transendothelial migration (**Figure 4B; Network 4**, Fisher’s exact test, Benjamini-Hochberg FDR < 0.1). Some notable associations for these shared host genes include host genes and taxa previously implicated in CRC and IBD. For example, expression of Annexin A1 or *ANXA1, a* host gene known to regulate intestinal mucosal injury and repair and found dysregulated in CRC and IBD ^133–135^, is positively correlated with Bacteroidales in CRC, while negatively correlated with Peptostreptococcaceae in IBD (**Figure 4B; Network 4**). Bacteroidales species are known to modulate maturation of the host immune system and gut barrier integrity ^136,137^. *TLR4*, a host gene known to modulate inflammatory response in intestinal epithelium through recognition of bacterial lipopolysaccharide ^138,139^, and previously implicated in IBD and CRC ^140,141^, is found associated with an oral microbe *GN02* in CRC ^142^, whereas in IBD, it interacts with Acidaminococcaceae, a gut microbe found increased in abundance in patients with Crohn’s disease ^143^ (**Figure 4B; Network 4**). Overall, our analysis shows that shared gut microbial taxa and shared host genes depict disease-specific host-microbe crosstalk, thus suggesting that the mechanism of host gene-microbiome interaction might be specific to the disease.

## Discussion

While gut microbial communities and host gene expression have separately been implicated with human health and disease, the role of the interaction between gut microbes and host gene regulation in the pathogenesis of human gastrointestinal diseases remains largely unknown. Here, we comprehensively characterized interactions between gut microbiome composition and host gene expression from 416 colonic mucosal samples taken from patients with colorectal cancer, inflammatory bowel disease, and irritable bowel syndrome, in addition to non-disease controls. To overcome the challenges associated with integrating high-dimensional multi-omic datasets, we developed and applied a machine learning framework to characterize interactions between the gut microbiome and host transcriptome in each disease. We identified a common set of host genes and pathways, including pathways that regulate gastrointestinal inflammation, gut barrier protection, and energy metabolism, that are associated with gut microbiome composition in all three diseases. We also found that gut microbes that have been previously associated with all three diseases, including *Streptococcus*, interact with different host pathways in each disease. This suggests that both common and disease-specific interplay between gut microbes and host gene regulation may contribute to the underlying pathophysiology of GI disorders.

Previous studies have found common microbial signatures across CRC, IBD, and IBS. For example, all three diseases exhibit an overrepresentation of Peptostreptococcaceae and *Streptococcus* spp ^8,43,62,95^. In addition, both CRC and IBD microbiomes are denoted by a loss of butyrate producing gut bacteria, including *Blautia*, and an enrichment of enterotoxigenic *Bacteroides fragilis* ^5,58,95,144^. In contrast to these microbiome similarities, host gene regulation shows distinct alterations across the three GI disorders; for example, unique antibacterial gene expression profile and disruption of purine salvage pathway are specific to IBS, deregulation of proinflammatory IL-23-IL-17 signaling is unique to IBD, and prominent activation of oncogenic pathways like Notch and WNT signaling is a hallmark of CRC ^8,13,145,146^. Here, we found that common disease-related gut microbes can interact with host genes and pathways in a disease-specific manner. Thus, it is compelling to hypothesize that although diseases can be characterized by similar microbial perturbations, these microbes can impact different pathophysiological processes through interaction with different host genes in each disease. For example, we found that, in CRC, *Streptococcus* is correlated with the expression of host genes that regulate WNT signaling and NF-κB pathway, whereas in IBS, *Streptococcus* is correlated with host genes that modulate macrophage inflammatory response, thus suggesting that this gut microbe may perturb distinct host pathways in CRC and IBS. Of course, since our results are based on correlational analysis, it is challenging to assess directionality. While it is possible that these disease-specific interactions have a role in disease pathogenesis, it is also possible that the disease-transformed colonic mucosa renders it more conducive to the same microbial taxa.

We also identified a common set of host genes and pathways that are associated with gut microbiome composition in all three diseases. These included pathways that regulate gastrointestinal inflammation, immune response, and energy metabolism, and have been previously implicated in these diseases ^33,147–149^. Our analysis shows that these common host genes and pathways correlate with disease-specific gut microbes in CRC, IBD, and IBS. For example, the expression of host gene *PINK1* that regulates the PI3-kinase/AKT signaling pathway ^126^ is associated with the abundance of *Collinsella* in CRC, Peptostreptococcaceae in IBD, and *Blautia* in IBS. This suggests that in some cases, distinct gut microbes may modulate host genes and pathways that are commonly dysregulated across different gut pathologies. At the same time, we also found disease-specific host gene-microbe interactions. For example, in CRC, Syndecan-1 pathway, a host pathway that modulates tumor growth and progression, is correlated with microbial taxa such as *Parvimonas* and *Bacteroides fragilis* that are known to promote intestinal carcinogenesis ^53,56,57,63^. These associations are not found in IBD or IBS, and are unique to CRC. Taken together, our results indicate that GI disorders are characterized by a complex network of interactions between microbes and host genes. Although these interactions can be disease-specific, we find cases where the same microbial taxon is associated with different host genes in each disease, and vice-versa: cases where the same host pathway is associated with different microbes in each disease. Although much effort in microbiome research has been directed towards identifying specific microbial taxa that are responsible for the pathogenesis of disease, our findings indicate that without incorporating data on host gene-microbe interactions, studies may be missing the full picture. Host omic data provides invaluable information on the potential mechanisms through which microbes can affect health.

An important contribution of our work is a machine learning-based integrative framework for characterization of complex host gene-microbe interactions across human diseases. Although few recent studies have investigated associations between host transcriptome and gut microbiome in human gut disorders, our analysis uses an innovative analytical technique that has several advantages ^24–27^. First, as opposed to analyses that rely on calculating pairwise correlations between features (e.g. Dayama et al.), our approach does not require restricting the data to a predetermined subset of taxa or genes of interest to increase statistical power. In addition, compared to Procrustes analysis, which is commonly used for finding overall correspondence between paired datasets, our approach does not only detect overall association, but can also find specific associations between gut microbial taxa and host genes (using Lasso) and pathways (using sparse CCA), allowing identification of specific interactions and shedding light on potential biological mechanisms of interaction. Furthermore, our approach can be applied to other types of multi-omic dataset, including microbial metabolomic and metagenomic data ^8^. Lastly, our project incorporates data from across several diseases, identifying commonalities across conditions as well as disease-specific patterns.

Despite these advantages, our study has several limitations. While we report the potential role of host gene-microbiome interactions in the pathophysiology of GI disorders, our study identifies correlations, and we cannot infer causality here. Given the challenges associated with studying causal mechanisms in humans, future studies using cell culture or animal models would be useful in elucidating the causal role and directionality of interactions between the gut microbiome and host gene regulation in these diseases ^150^. Another caveat of our study is that it includes three different disease cohorts with disparate sample collection and sequencing protocols, which can lead to potential batch effects. To mitigate this issue, we performed our integration analysis in each disease cohort separately, including cases and internal controls within each cohort. Additionally, our analysis focused only on the taxonomic composition of the microbiome, and hence we could not characterize interactions involving microbial genes and pathways. Lastly, there are several environmental variables that could potentially influence the microbiome, including diet and medication history, which are not available across our disease cohorts.

Overall, our work demonstrates the power of integrating gut microbiome and host gene expression data to provide insights into their combined role in GI diseases, including CRC, IBD, and IBS. We find disease-specific and shared gut microbe-host gene interactions across these gut disorders, involving gut microbes and host genes implicated in gastrointestinal inflammation, gut barrier protection, and metabolic functions. We also found that the same gut microbes interact with different host genes in different diseases, suggesting potential mechanisms by which similar gut microbes can affect different disease pathologies. These results represent an important step towards characterizing the crosstalk between gut microbiome and host gene regulation and understanding the contribution to disease etiology.

## Acknowledgements

We would like to thank the IBD HMP2 consortium for making the dataset publicly available. We are thankful to the Blekhman Lab members for their comments and suggestions on the manuscript. We thank Dr. Wen Wang and Dr. Gabriel Al-Ghalith for their feedback. This work is supported by NIH grant R35-GM128716 (to R.B.), a University of Minnesota Doctoral Dissertation Fellowship (to S.P), and by NIH grant R01-GM130622 (to E.F.L). This work was carried out, in part, by resources provided by the Minnesota Supercomputing Institute.

## Methods

### Overall study design, samples and data

We obtained mucosal microbiome (16S rRNA) and host gene expression (RNA-seq) data from colonic mucosal biopsy samples collected from the patients from three disease cohorts: colorectal cancer (CRC), inflammatory bowel disease (IBD), and irritable bowel syndrome (IBS). Except the host gene expression (RNA-seq) data for CRC, all the other datasets were generated and described in detail in previous studies ^3,8,25^. Below, we describe the sample collection, sequencing, and quality control for host RNA-seq data for CRC cohort, and summarize data acquisition process for other datasets:

### CRC samples and data

We used 88 pairs of colonic mucosal samples from 44 patients, with primary tumor and normal tissue samples from each individual. These samples were characterized and described in a previous study ^3^. Detailed cohort characteristics are included in Supplementary Table S1.

#### Host RNA-seq sequencing, alignment and quality control

Total RNA was extracted using a previously established protocol ^3,151^. Approximately 100mg of flash-frozen tissue per sample were lysed by placing the tissue in 1mL of Qiazol lysis reagent (Qiagen Inc., Valencia, CA, USA) and sonicating in a 65° C water bath for 1-2 hours. Nucleic acids were purified from the lysates using the Qiagen AllPrep DNA/RNA mini kit (Qiagen Inc., Valencia, CA, USA), quantified using a Nanodrop 2000 spectrophotometer (Thermo Fisher Scientific, Waltham, MA USA), and submitted for RNA sequencing to the University of Minnesota Genomics Center. Total eukaryotic RNA isolates were quantified using a fluorimetric RiboGreen assay, and once the samples passed the initial QC step (≥ 1 microgram and RIN ≥ 8), they were converted to Illumina sequencing libraries using Illumina’s TruSeq Stranded Total RNA Library Prep (for details, see https://www.illumina.com). Truseq libraries were hybridized to a paired-end flow cell and individual fragments were clonally amplified by bridge amplification on the Illumina cBot. Once clustering was complete, the flow cell was loaded on the HiSeq 2500 and sequenced using Illumina’s SBS chemistry. Base call (.bcl) files for each cycle of sequencing were generated by Illumina Real Time Analysis (RTA) software. Primary analysis and index de-multiplexing are performed using Illumina’s bcl2fastq v2.20.0.422, which output the demultiplexed FASTQ files.

A quality check of raw sequence FASTQ files was performed using FastQC software (version 0.11.5) ^152^. Quality trimming was performed to remove sequence adaptors and low quality bases using Trimmomatic with 3bp sliding window trimming from 3’ end requiring minimum Q16 (phred33) ^153^. FastQC was run on the resulting trimmed files to ensure good quality of sequences. The paired-end reads were mapped to NCBI v38 *H. sapiens* reference genome using HISAT2 ^154^, resulting in an average alignment rate of 87.11% overall for 88 samples. We obtained a range of read counts between 14,365,657 and 31,530,487 aligned reads per sample, with an average of 22,475,688.2 and 22,697,605.5 aligned reads per sample. SAMtools was used for sorting and indexing the aligned bam files. After alignment, the *Subread* package (version 1.4.6) within the *featureCounts* program was used to generate transcript abundance file ^155^ (Supplementary Figure S4).

#### 16S rRNA data acquisition

The microbiome dataset used in this study was generated and characterized previously ^3^. We used the unnormalized and unfiltered OTU table in tab-delimited format, representing mucosal microbiome data from 44 tumor and 44 patient-matched colon tissue samples.

### IBD samples and data

We used previously generated and described host gene expression (RNA-seq) and mucosal gut microbiome (16S rRNA) data for the IBD cohort generated as part of the HMP2 project ^25,28^ (for detailed protocols, see http://ibdmdb.org/protocols). These include data from colonic biopsy samples collected from 78 individuals, including 56 individuals with IBD, and 22 individuals without IBD (“non-IBD” in HMP2). Out of 56 IBD patients, 34 patients had Crohn’s disease (CD) and 22 patients had ulcerative colitis (UC). Detailed cohort characteristics are included in Supplementary Table S1. We downloaded metadata, host RNA-seq data, and microbiome data for these samples from http://ibdmdb.org in July 2018. We downloaded the unnormalized and unfiltered OTU table and host transcript read counts files in tab-delimited format. We describe the filtering and preprocessing steps for host gene expression and microbiome data below.

### IBS samples and data

We used previously generated and characterized host gene expression (RNA-seq) and mucosal gut microbiome (16S rRNA) data for the IBS cohort ^8^. These include data from colonic biopsy samples collected from 42 individuals, including 29 individuals with IBS, and 13 healthy individuals (non-IBS). Detailed cohort characteristics are included in Supplementary Table S1. We obtained the unnormalized and unfiltered OTU table and host transcript read count files in tab-delimited format via personal communication with authors of the paper ^8^. For some individuals, samples were collected at two time points. For these cases, we averaged the gene expression levels and microbiome abundance measurements across the two time points. This is supported by a recent study showing that “omics” methods are more accurate when using averages over multiple sampling time points ^156^. We describe the filtering and preprocessing steps for host gene expression and microbiome data below.

### Preprocessing host gene expression data

For host gene expression data for each disease cohort, we used *biomaRt* R package (version 2.37.4) to only keep data for protein-coding genes ^157^. We filtered out low expressed genes to retain genes that are expressed in at least half of the samples in each disease cohort. We performed variance stabilizing transformation using the R package *DESeq2* (version 1.14.1) on the filtered gene expression read count data ^158^. We filtered out genes with low variance, using 25% quantile of variance across samples in each disease cohort as cutoff. Performing these steps for RNA-seq data for each disease cohort separately resulted in a unique host gene expression matrix per disease for downstream analysis, including 12513 genes in the CRC dataset, 11985 genes in IBD dataset, and 12429 genes in IBS dataset.

### Preprocessing microbiome data

We performed the following steps for microbiome data from each disease cohort separately. First, sequences that were classified as either having originated from archaea, chloroplasts, known contaminants originating from laboratory reagents or kits, and soil or water-associated environmental contaminants were removed from the OTU table as described previously^159^. Next, we summarized the OTU table at the species (if present), genus, family, order, class, and phylum taxonomic levels, and performed prevalence and abundance-based filtering to retain taxa found at 0.001 relative abundance in at least 10% of the samples. We then concatenated these summarized taxa matrices (count data) into one combined taxa matrix for each disease dataset. We applied centered log ratio (CLR) transform on the filtered taxa count matrix to account for compositionality effects. These steps resulted in a taxonomic abundance matrix for each disease cohort, which included 235 taxa in the CRC dataset, 121 taxa in the IBD dataset, and 238 taxa in the IBS dataset.

### Procrustes analysis

To assess overall correspondence between host gene regulation and gut microbiome composition in CRC, IBD, and IBS, we performed Procrustes analysis in R using the *vegan* package (version 2.4-5) ^160^. For each disease cohort, we used Aitchison’s distance on host gene expression data and Bray Curtis distance on gut microbiome data as input to the Procrustes analysis ^161^. The significance of rotation agreement was obtained using the *protest()* function with 9,999 permutations.

### Sparse Canonical Correlation Analysis

We used sparse canonical correlation analysis (sparse CCA) to identify group-level correlations between paired host gene expression and gut microbiome data in each disease cohort. Canonical correlation analysis (CCA) identifies linear projection of two sets of observations into shared latent space that maximizes correlation between the two datasets ^162^. Sparse CCA is adapted from CCA for high dimensional settings to incorporate feature selection by utilizing *L1* or lasso penalty in CCA ^29^. The objective function of sparse CCA can be expressed as follows:

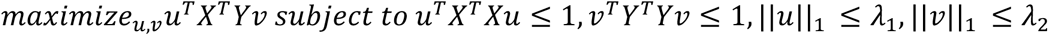
where, *X* and *Y* denote two data matrices with same number of samples, but different number of features (representing gut microbiome taxonomic composition data and host gene expression data, respectively); *u* and *v* are canonical loading vectors of X and Y respectively; *λ* and *λ*_2_ control lasso penalties of *u* and *v*, respectively.

For each disease cohort separately, we applied sparse CCA using R (version 3.3.3) package *PMA* (version 1.1) with gut microbiome taxonomic composition and host gene expression as two sets of variables to be correlated ^163^. Below, we describe details on hyperparameter tuning, fitting sparse CCA models, computing significance of correlation for sparse CCA components, enrichment analysis, and visualization of sparse CCA output.

### Hyperparameter tuning and fitting for Sparse CCA model

We performed hyperparameter tuning to identify the sparsity penalty parameters for gut microbiome abundance (λ_1_) and host gene expression (λ_2_) data. Since the permutation search provided in the *PMA* package only performs a one-dimensional search in the tuning parameter space, we implemented a grid-search approach using leave-one-out cross-validation in R (version 3.3.3) for hyperparameter tuning. We selected penalty parameters which had the highest correlation under cross-validation. Using this approach, we identified λ_1_ as 0.15 and λ_2_ as 0.2 for CRC data, λ_1_ as 0.177 and λ_2_ as 0.333 for IBD data, λ_1_ as 0.4 and λ_2_ as 0.1 for IBS data.

After identifying sparsity parameters, we fit the sparse CCA model to obtain subsets of correlated host genes and gut microbes, known as components. Each sparse CCA component includes non-zero weights (or canonical loadings) on gut microbes, and non-zero weights on a subset of host genes correlated with those gut microbes to capture joint variation in the two sets of observations. We computed the first 10 sparse CCA components for each disease cohort, performing a separate computation for case and control samples. Sparse CCA components are computed iteratively, informed by previously computed components, thus, resulting in uncorrelated components ^164^. Next, we assessed the significance of sparse CCA components as described below.

### Significance of correlation for sparse CCA components

We computed the significance of each pair of canonical variables (or a component) using leave-one-out cross-validation approach in R (version 3.3.3). For a given component, we first used the penalty parameters determined above to compute the sparse CCA output with one sample held out. We then computed the scores for the held-out sample, i.e., we computed *scoreX_i_ = X_i_u_−i_* and *scoreY_i_ = Y_i_v_−i_*, where *i* is the held-out sample, *X_i_*and *Y_i_* denote the values for *i^th^* sample of the input data matrices *X* and *Y*, and *u_−i_* and *v_−i_* are the canonical loadings estimated from the sparse CCA computation without the *i^th^*sample. We repeated this *n* times, where *n* is the total number of samples in the data, to obtain the vector of held-out scores. To assess the true strength of association and its significance, we used *cor.test()* on the scores computed for the held-out samples. We corrected the p-values for multiple hypothesis testing using Benjamini-Hochberg (FDR) method within each disease cohort, and determined significant components at FDR < 0.1.

Using this approach, we identified 7 significant components in CRC, with an average of 828 host genes and 8 gut microbes; 4 significant components in IBD, with an average of 2095 host genes and 6 gut microbes; and 6 significant components in IBS, with an average of 577 host genes and 61 gut microbes (FDR < 0.1, Supplementary Tables S2-S4).

### Enrichment analysis for sparse CCA

To characterize host pathways enriched for the set of host genes associated with microbes in each component, we implemented an enrichment analysis in R (version 3.3.3). We implemented Fisher’s exact test to assess pathway enrichment, where we used the set of host genes input to the sparse CCA analysis as background genes, and set of host genes in a component as the genes of interest. We used KEGG and PID gene sets from MsigDB canonical pathways collection ^165,166^. To avoid pathways that are too large to provide any specific biological insights or too small to provide adequate statistical power, we excluded any pathway with either (1) fewer than 25 genes, (2) more than 300 genes, or (3) fewer than 5 genes that overlapped between the genes of interest and the pathway. We combined the set of enriched host pathways for all significant components for a given disease dataset, corrected for multiple hypothesis testing within each disease cohort using Benjamini-Hochberg (FDR) approach, and determined significant host pathways at FDR < 0.1. This analysis was performed separately for case and control data for each disease.

To identify case-specific host pathways, we used a two-part approach: (1) first, we identified pathways that are only significantly enriched in cases (FDR < 0.1) and not in controls. (2) In addition, we identified pathways that are significant in both the cases and controls at FDR < 0.1. For these pathways, we performed differential enrichment in cases versus controls by implementing a comparative log odds-ratio approach in R ^167,168^. To do so, we first computed the z-score for the odds ratio for *i*-th pathway in cases:

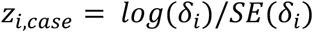

where, *δ_i_*, is the odds-ratio for *i*-th pathway in cases, and SE (*δ_i_*) is the standard error for *i*-th pathway in cases, which is computed using the four elements, n_1_to n_4_, of the 2×2 contingency table used in the enrichment analysis for the *i*-th pathway as follows:

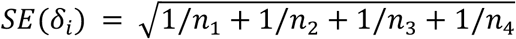

Similarly, we computed *Z_i,ctrl_* for the same pathway in the controls. For a given pathway, we compare enrichment for the component that gives highest significance for the cases with that which gives highest significance for the controls. Next, we compute a comparative log odds-ratio for *i*-th pathway overlapping between cases and controls as follows:

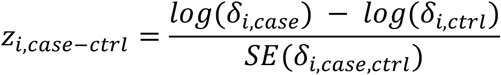

The greater the value of *Z_i,case−ctrl_*, the greater the odds a pathway is differentially enriched in case versus control than by chance. P-values were inferred assuming normal approximations, and corrected for multiple hypothesis testing using Benjamini-Hochberg (FDR) approach. As the last step of part (2), we retained pathways that were differentially enriched in cases versus controls at FDR < 0.2. Finally, we combined the pathways from part (1) and (2) to obtain case-specific pathways.

### Visualizing disease-specific and shared host pathways and components from sparse CCA

To determine *shared* host pathways, i.e. host pathways for which gene expression correlates with gut microbes across all three disease cohorts, and *disease-specific* host pathways, i.e. host pathways for which gene expression correlates with gut microbes in only one of the three disease cohorts, we computed overlaps between significant case-specific host pathways determined above across the three disease cohorts. Given the overlap across the curated gene sets from MsigDB, we controlled for redundancy across pathways for visualization purposes. To do this, we identified similar pathways based on their relative overlap in terms of the set of genes using an overlap coefficient. The overlap coefficient between two pathways is defined as the number of common genes between the pathways divided by the number of genes in the pathway with fewer genes. Specifically, the overlap coefficient is represented as follows:

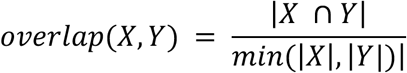

For the top 15 most significant host pathways (FDR < 0.1) discovered for each shared and disease-specific set (Supplementary Table S12), we computed pairwise similarity between pathways as overlap coefficients and used a maximum allowed similarity score of 0.5 as a cutoff. Using the pairs of pathways that satisfied the cutoff, we computed the connected components to identify clusters of overlapping pathways. For visualization purposes, we selected a representative pathway from each connected component, prioritising the pathway with the highest number of genes (Figure 2A, Supplementary Table S4). We visualized host pathway enrichment using the R package *ggplot* (version 3.2.1).

For visualizing components corresponding to selected host pathways or common host genes across diseases (Figures 2B - D), we ordered host genes and taxa by their absolute coefficients in the component, and selected the top 10 host genes and taxa for representation. If multiple taxa originating from the same lineage occurred in a component, we selected the one with the highest coefficient to reduce redundancy, thus representing the taxa with most contribution from a given lineage. The size of host genes and gut microbial taxa are scaled by the absolute value of their corresponding coefficients in a given component. All sparse CCA components were visualized using Cytoscape (version 3.5.1) ^169^.

### Lasso regression analysis

We used Lasso penalized regression to identify specific interactions between individual host genes and gut microbial taxa within each disease cohort ^31^. We implemented a gene-wise model using expression for each host gene as response and abundances of microbiome taxa as predictors, to identify microbial taxa that are correlated with a host gene. An ordinary least squares (OLS) regression is not suitable for this task, since OLS results in unstable solutions under high-dimensional settings or when p >> *n*, i.e. number of predictors p is much larger than number of samples *n.* Additionally, we expect the abundance of only a few microbial taxa to correlate with the expression of each host gene. To address this, we used lasso regression, which is similar to multivariate OLS, except that it uses shrinkage or regularization to perform variable selection, thus picking only a few taxa that associate with a host gene’s expression.

To account for other factors that can impact host gene expression or microbiome composition, each model also included covariates in the predictor matrix (i.e. microbiome abundance table) for gender (male or female), disease-subtype for IBD (Crohn’s Disease or ulcerative colitis), disease-subtype for IBS (constipation (IBS-C) or diarrhea (IBS-D)).

The lasso model estimates the lasso regression coefficient 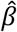 by minimizing the following:

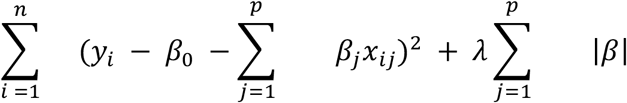
where, *n* = number of samples; *p* = number of predictors (taxa and other covariates); 1 ≤ *i* ≤ *n*, 1 ≤ *j* ≤ p; *y* = response (host gene expression); *x* = predictor (taxa abundance and other covariates); *λ* = tuning parameter, *λ* ≥ 0.

In addition to minimizing the residual sum of squares (first term in the equation), lasso minimizes the *l*_1_ norm of the coefficients (second term in the equation), which has an effect of forcing some of the coefficients to zero as the value of tuning parameter,*λ*, increases. Thus, lasso performs feature or variable selection that leads to sparse models.

We implemented a lasso regression framework using R (version 3.3.3) package glmnet (version 2.0-13), which uses cyclical coordinate descent to compute regularization path ^170^. Our framework executes a lasso regression for each host gene’s expression as response and abundances of microbial taxa and values of other covariates as predictors. We used leave-one-out cross-validation to estimate the tuning parameter, *λ*, which was used to fit the final model on a given disease dataset.

We then performed inference for the lasso model using a regularized projection approach known as desparsified lasso. The desparsified lasso uses the asymptotic normality of a bias-corrected version of the lasso estimator to obtain 95% confidence intervals and p-values for the coefficient of each predictor (microbe) associated with a given host gene ^171^. We used the R package *hdi* (version 0.1-7) that implements the desparsified lasso approach for estimation of confidence intervals and hypothesis testing in high dimensional and sparse settings ^171,172^. We then corrected for multiple hypothesis testing using Benjamini-Hochberg (FDR) method.

### Stability selection for Lasso model

Since the lasso model is sensitive to small variations of the predictor variable, we used stability selection to pick out robust microbes associated with a host gene ^173^. Stability selection is a resampling-based method that can be combined with different variable selection procedures in high dimensional settings, including lasso. Briefly, stability selection with lasso proceeds as follows:

Step 1. Select a random subset of the data.
Step 2. Fit the lasso model with a randomly perturbed penalty term in the neighborhood of the “best” penalty *λ*. Record the set of selected variables (microbes).
Step 3. Repeat steps 1) and 2) *K*times.
Step 4. Compute the frequency of selection, *f_i_* per variable (microbe) across all trials.
Step 5. Select the variables (microbes) that are selected with a frequency of at least *f_thr_*, a pre-specified threshold value. Thus, we select a set of stable variables (microbes) such that *f_i_* ≥ *f_thr_*.

The overall idea is that, if the same variables (microbes) are repeatedly selected when the parameters are perturbed, then they are robust variables. Stability selection also controls for family-wise error rate, thus controlling for false positives in addition to the FDR approach mentioned above ^173^. In our analysis, we used the R package *stabs* (version 0.6-3) to perform stability selection ^174^. Specifically, we used the following parameters in the process described above: in Step 1, a random subset of size *n/2* of data is selected, where nis total number of samples, in Step 3, *K* = 100, and in Step 5, *f_thr_* = 0.6, i.e. a predictor (microbe) selected in at least 60% of the fitted models is considered stable. The choice of these parameters are in accordance with the proposal of stability selection by Meinshausen and Bühlmann^173^.

Finally, we performed an intersection between associations identified by stability selection here and associations identified at FDR < 0.1 by the lasso model described above. We removed any host gene-gender and host gene-disease-subtype associations to obtain the significant and stability selected host gene-microbe associations at FDR < 0.1.

### Parallel execution of lasso analysis on supercomputing nodes

We implemented a parallel framework for executing the gene-wise lasso analysis, where we parallelized execution of lasso models on host genes across multiple nodes and cores on a compute cluster from Minnesota Supercomputing Institute. We used job arrays to parallelize our analysis on multiple nodes on the cluster. Additionally, we used R packages *doParallel* (version 1.0.15) and *foreach* (version 1.4.7) to run parallel processes on multiple cores of each compute node.

### Enrichment analysis for lasso output

To characterize biological functions for the host genes that were found associated with specific gut microbes in a disease cohort by the lasso framework, we implemented an enrichment analysis in R (version 3.3.3) using Fisher’s exact test. We used the set of expressed genes input to the lasso analysis as the background genes, and the set of host genes associated with gut microbes in a patient samples as genes of interest. We used the KEGG, PID, and REACTOME gene sets from MsigDB canonical pathways collection ^165,166^. To avoid too large or too small pathways, we excluded from our analysis any pathway with fewer than 25 genes, greater than 85 genes, or fewer than 5 genes that overlap between the genes of interest and the pathway. The p-values obtained from Fisher’s exact test were adjusted for multiple testing using Benjamini-Hochberg (FDR) approach. We identified 87 host pathways that are unique to each disease, including 22 CRC-specific, 60 IBD-specific, and 5 IBS-specific pathways that interact with unique gut bacteria (FDR < 0.2, Supplementary Table S8). Here, we used a more relaxed FDR threshold of 0.2 to present a larger number of biologically relevant host pathways.

### Visualization of shared genes and taxa interactions for lasso output

In Figure 4, we visualized host gene-microbe interactions for gut microbes and host genes shared across diseases. For visualizing interactions for shared gut microbes (Figure 4A), we identified shared microbes between all possible overlaps between diseases (Figure 4A; Networks 1–4), host genes that interact with these common microbes in each disease, and all interactions involving these microbes and genes in each disease (FDR < 0.1). Next, we grouped gene-taxa interactions identified per disease by shared taxa, sorted them by FDR value, and picked top gene-taxa association per shared taxa until we obtained at most 10 interactions per disease (FDR < 0.1, Supplementary Table S9).

Similarly, for visualizing interactions for shared host genes (Figure 4B), we identified shared host genes between all possible overlaps between diseases (Figure 4B; Networks 1–4), and host gene-taxa interactions per disease for these host genes shared across diseases. We sorted the interactions by FDR adjusted q-values (ordered first by q-value in CRC associations, followed by q-value in IBD associations, and finally by q-value in IBS associations, depending on the overlapping set under consideration). We picked top 10 genes from this merged output, and identified at most top 10 associations involving these genes in each disease for the overlapping set under consideration (FDR < 0.1, Supplementary Table S10). Since lasso gives biased estimates of the coefficients, we used Spearman correlation coefficient (rho) to depict strength of association for visualizing host gene-taxa associations. All the associations in Figure 4 were visualized using Cytoscape v3.5.1, where shared features are in grey and disease-specific features in disease-specific colors ^169^.

## Data and Software Availability

Raw data for host RNA-seq for CRC cohort is available on the NCBI Sequence Read Archive (SRA) under submission ID: SUB9143781. For raw data from 16S rRNA sequencing for CRC cohort, RNA-seq and 16S rRNA sequencing for IBD cohort, and RNA-seq and 16S rRNA sequencing for IBS cohort, please refer to data accession details published previously ^8,25,28^. Processed data tables for host transcriptomics and microbiome data for each disease cohort have been included as supplemental tables (Supplementary Tables S13–S18). Code used for integration analyses performed in the paper is available at:

https://github.com/blekhmanlab/hostgenemicrobiomeinteractions

## Ethics statement

For the colorectal cancer cohort, all research conformed to the Helsinki Declaration and was approved by the University of Minnesota Institutional Review Board, protocol 1310E44403. For the inflammatory bowel disease and irritable bowel syndrome cohorts, ethical approval is described in their respective publications ^8,25,28^.

