## Supplementary figures for "Shared and disease-specific host gene-microbiome interactions across human diseases"

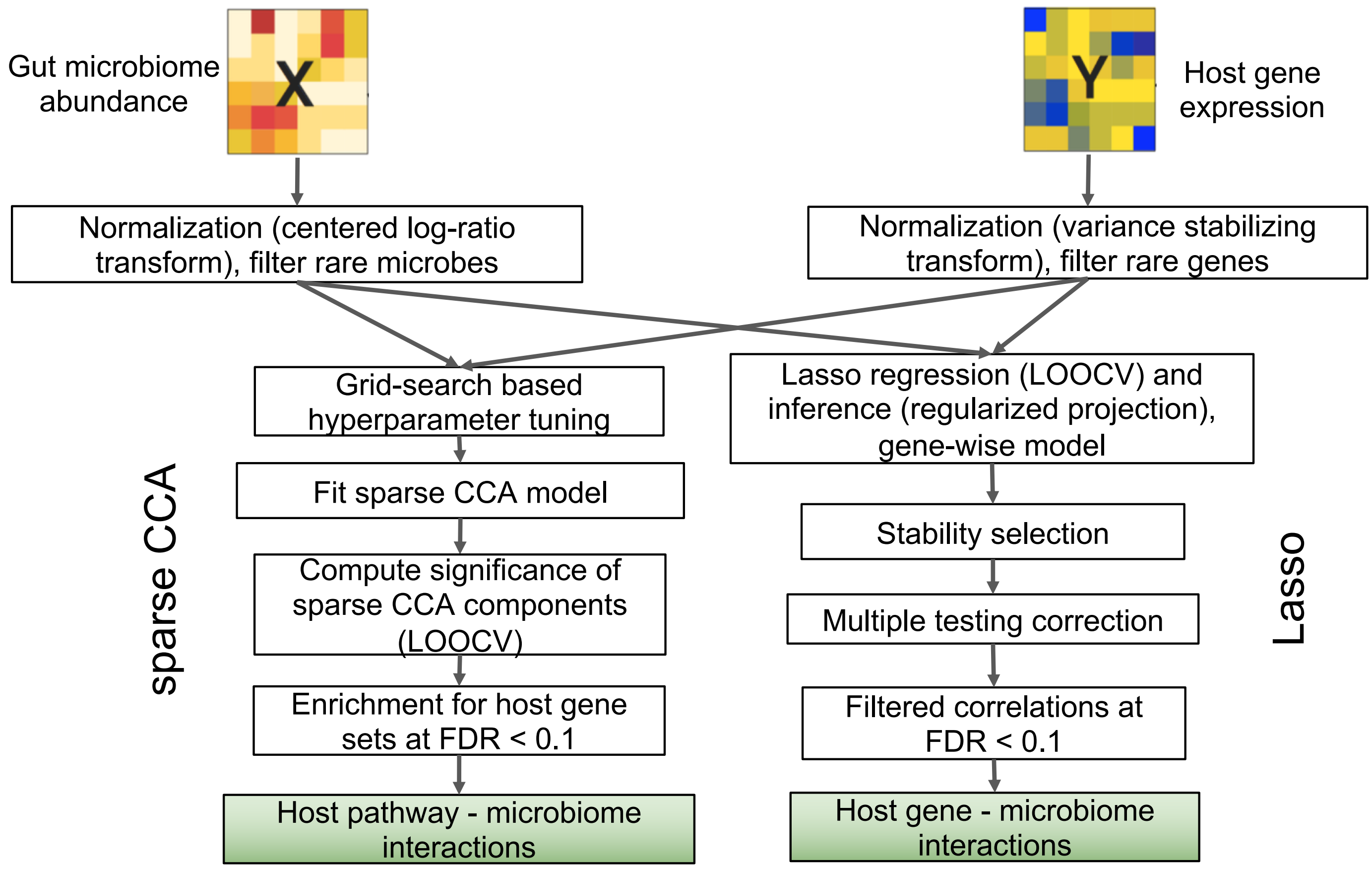


Figure S1: Overview of host gene-microbiome integration pipeline.

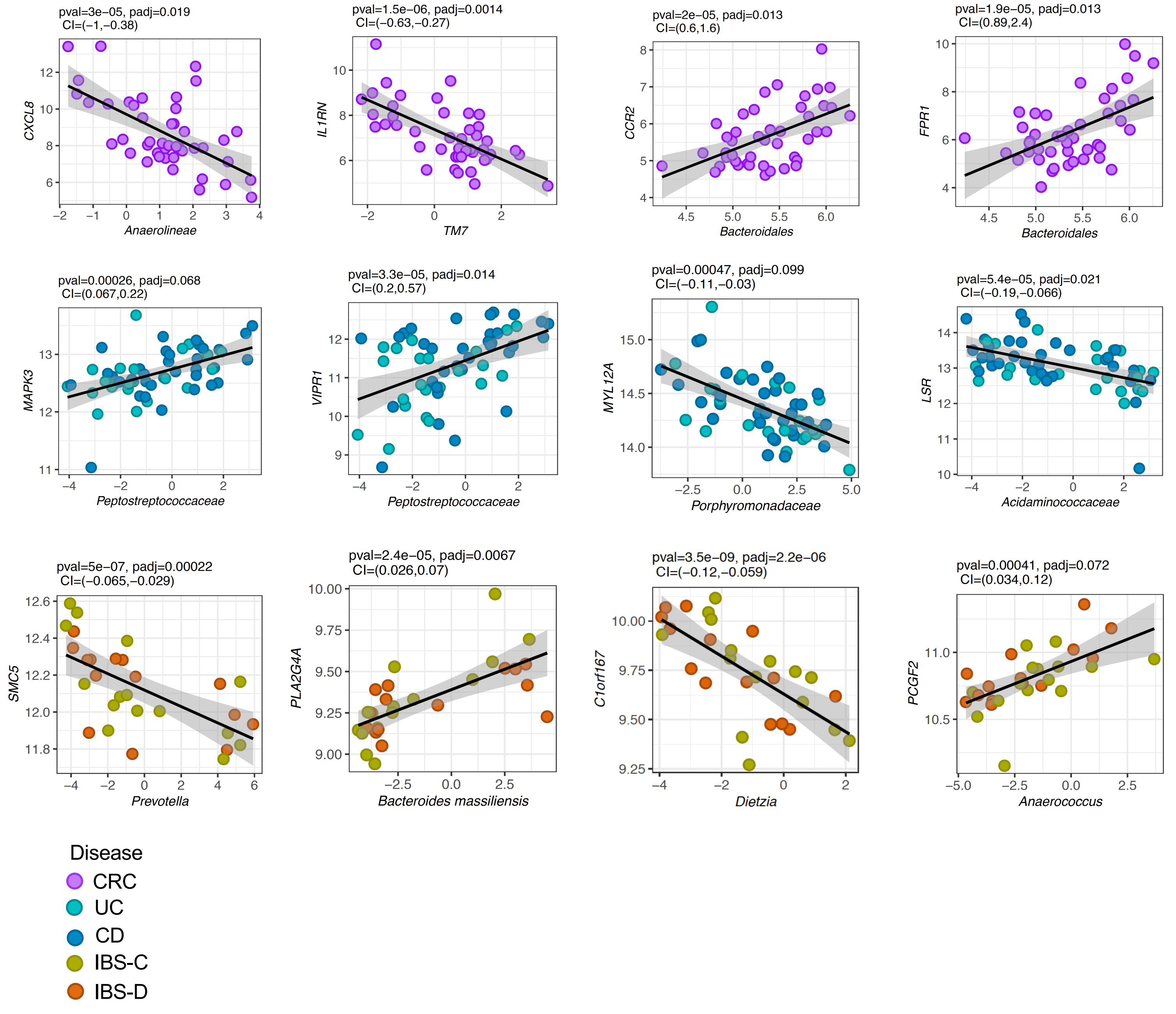


Figure S2: Examples of specific interactions between gut microbial taxa and host genes in CRC (top row), IBD (middle row), and IBS ( bottom row). The x-axis represents normalized abundance of microbial taxa, and the y-axis represents normalized expression of host gene. CRC: colorectal cancer, UC: Ulcerative colitis, CD: Crohn’s Diseases, IBS-C: Irritable bowel syndrome - constipation, IBS-D: Irritable bowel syndrome – diarrhea.

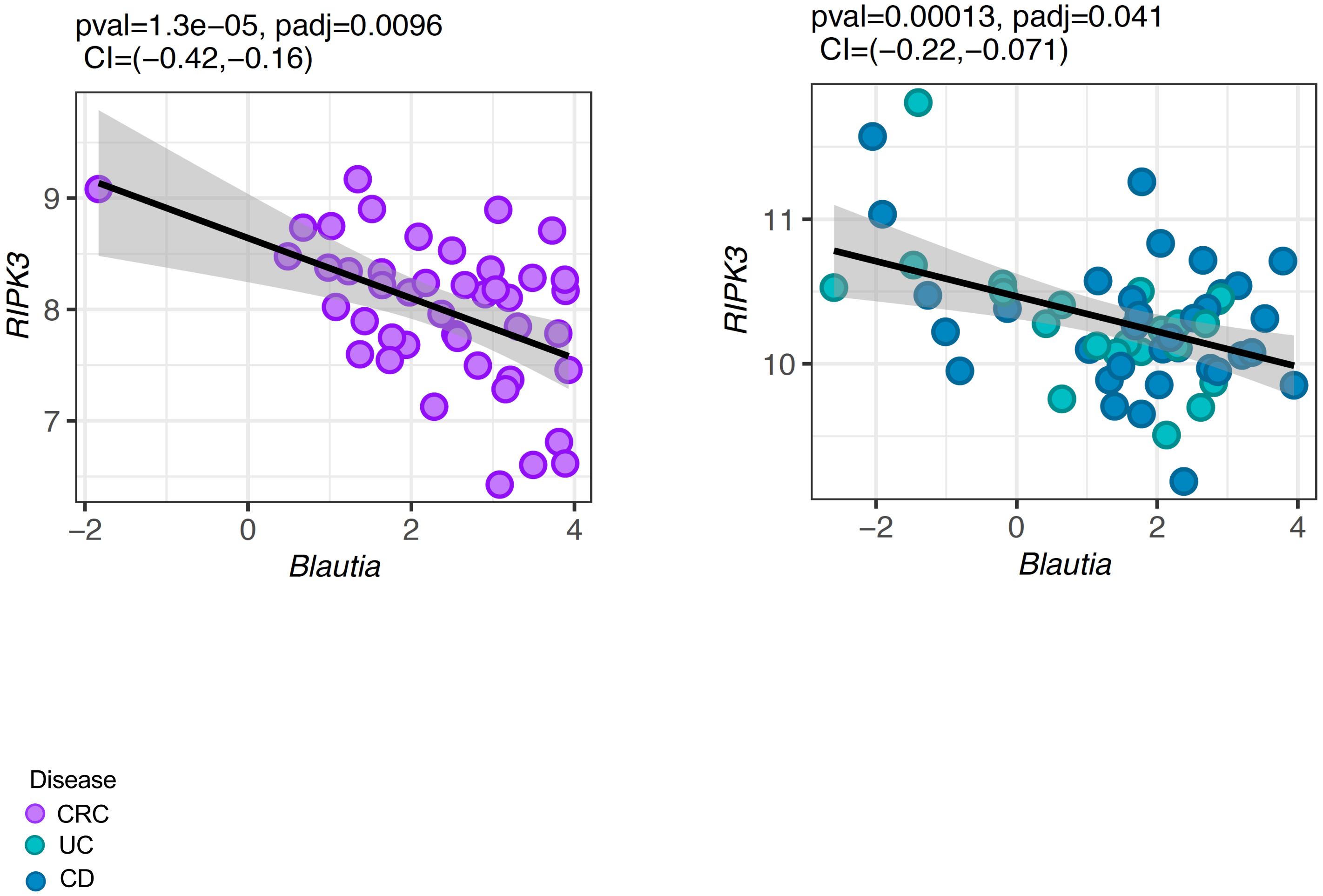


Figure S3: A common host gene-microbe interaction between CRC and IBD. CRC: colorectal cancer, UC: Ulcerative colitis, CD: Crohn's Diseases.

A

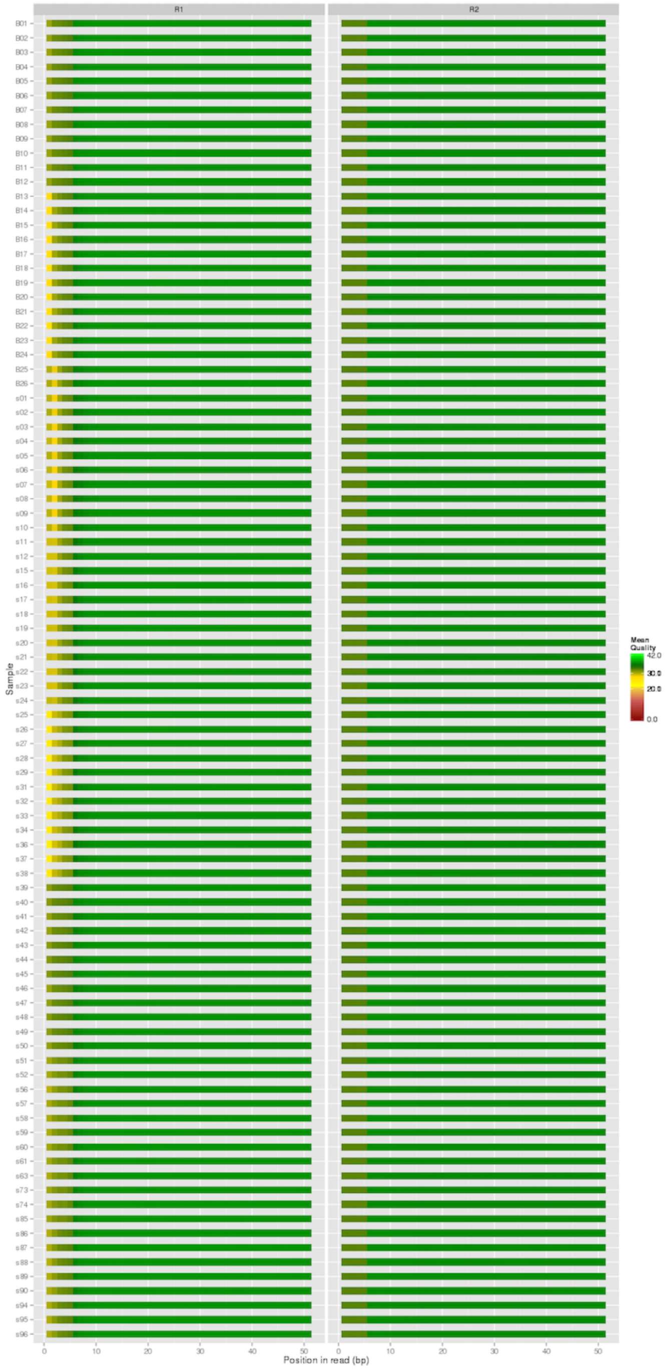


B

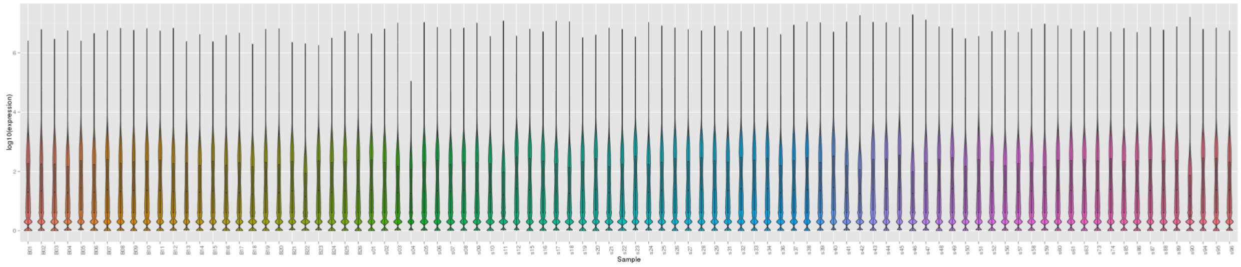


Figure S4: Quality control and transcript quantification of host RNA-seq data for CRC samples. A. Average phred score for forward (R1) and reverse (R2) reads output by FASTQC. B. Distribution of host gene expression ($log_{10}$(expression) value) for each sample quantified by *Subread* package.
